# A sample preparation protocol for high throughput immunofluorescence of suspension cells

**DOI:** 10.1101/2020.01.05.895201

**Authors:** Anna Bäckström, Laura Kugel, Christian Gnann, Hao Xu, Joseph E. Aslan, Emma Lundberg, Charlotte Stadler

## Abstract

Imaging is a powerful approach for studying protein expression and has the advantage over other methodologies in providing spatial information *in situ* at single cell level. Using immunofluorescence and confocal microscopy, detailed information of subcellular distribution of proteins can be obtained. While adherent cells of different tissue origin are relatively easy to prepare for imaging applications, nonadherent cells from hematopoietic origin, present a challenge due to their poor attachment to surfaces and subsequent loss of a substantial fraction of the cells. Still, these cell types represent an important part of the human proteome and express genes that are not expressed in adherent cell types. In the era of cell mapping efforts, overcoming the challenge with suspension cells for imaging applications would enable systematic profiling of hematopoietic cells. In this work, we successfully established an immunofluorescence protocol for preparation of suspension cell lines and peripheral blood mononucleated cells (PBMC) and human platelets. The protocol is based on a multi-well plate format with automated sample preparation, allowing for robust high throughput imaging applications. In combination with confocal microscopy, the protocol enables systematic exploration of protein localization to all major subcellular structures.

## Introduction

Immunofluorescence (IF) is an application widely used to study subcellular localization of proteins to gain knowledge of protein function and it’s interaction partners. Over the past fifteen years, our group has contributed to establish the Human protein Atlas (HPA) database, where immunohistochemistry and immunofluorescence has been applied to systematically explore protein expression in 44 different tissue types and subcellular protein localization in selected cell lines (1–3). The current version of the protein atlas database contains tissue profiles of nearly 20,000 proteins and subcellular localization for over 12,000 proteins. The ultimate goal of this initiative is to generate a complete map of the human proteome at both tissue and subcellular level. While most proteins are expressed and successfully stained in one or several of the 44 tissue types, detection of all proteins in cell lines as used for the subcellular profiling is a challenge. Studies using mass spectrometry show that most cell lines express between 10-12,000 genes (4, 5). Within the HPA, the transcriptome of a large number of cell lines has been analyzed using RNA sequencing (6) and the number of cell lines being used for immunofluorescence and subcellular profiling has gradually increased to detect more of the protein coding genes (7). The transcriptomics data from the 64 cell lines being sequenced, reveal that over 4000 genes show highest expression in any of the 16 cell lines from hematopoietic origin. Further, approximately 700 of those genes are only expressed within this group of cell lines, making them a highly valuable from a cell mapping perspective. However most of these cell lines are not ideally suited for imaging, since they normally grow in suspension and do not readily attach to substrates in a manner needed for imaging.

As a major obstacle with suspension cells is poor attachment, several coatings have been developed with the aim to better attach the cells to the surface (8, 9). Most coatings used today are a variety of biological materials including extracellular matrix (ECM) proteins such as collagen, laminin and fibronectin. Basic synthetic polymers, like poly-D-lysine (PDL) and poly-ornithin are also popular coatings. Some protocols and equipment have been developed trying to mechanically help cells stick to the surface (10, 11). Even if cells initially manage to attach to the plates or glass slides, the shear stress and flow from pipetting during the sample preparation (i.e. antibody staining) is in many cases brutal, and very few cells retain at the surface. A typical immunofluorescence protocol consists of several pipetting steps including fixation, permeabilization, primary and secondary antibody incubation and many washing steps in between. In many cases the sample preparation protocol itself cannot be compromised to achieve high quality images in the end. To minimize the pipetting of the cells on the slides, some immunofluorescence protocols are designed to perform most steps in solution to at the end of the protocol gentle centrifuge the cells down into the imaging plate (12). However, this is not ideal as the attachment of already fixed cells to a surface is more challenging as cells are stiffer and less flexible. Second, the benefit from observing interactions between cells are lost using this approach.

Although these efforts have resulted in better prerequisites for imaging of suspension cells, protocols for systematic high throughput and high-quality imaging of suspension cells are still lacking. Here, we present an automated immunofluorescence protocol for suspension cell lines and peripheral blood cells (i.e., PBMC and platelets), suitable for systematic protein profiling at high throughput.

## Materials and methods

### Cell cultivation and coatings

For testing of different coating substrates the T-cell derived JURKAT cell line (ACC 282, DSMZ Leibniz, Germany) was used. For testing and optimization of the automated sample preparations THP1 (ACC-16), HEL (ACC-11), K-562 (ACC-10), REH (ACC-22) all from DSMZ, Leibniz Germany, cells were used in addition to the Jurkat cells. The adherent osteosarcoma cell line U-2 OS (HTB-96, ATCC Wesel, Germany) was used as a reference for comparison of attachment. All cell lines were cultivated following recommendations from the providers and incubated at 37 °C and 5.2 % CO_2_. For immunofluorescence 96 well glass bottom plates (Greiner Sensoplate 655891, BioNordica Stockholm, Sweden) were coated with fibronectin (FN) (VWR, 734-0101), poly-L-ornithine (PLO) (Sigma-Aldrich, A-004-M), poly-L-lysine (PLL) (Sigma-Aldrich, P8920) and laminin (LN) (Sigma-Aldrich, L2020) at different concentrations as shown in Error! Reference source not found.

The plates were either incubated with the respective FN or LN coating solution for 1 hour at RT (coating solution was removed afterwards) or incubated with PLO or PLL coating solution for 5 minutes at RT followed by a washing step with 1x phosphate buffered saline (1x PBS; 137 mM NaCl, 2.7 mM KCl, 10 mM Na_2_HPO_4_, 1.8 mM KH_2_PO_4_). Also Matrigel (catalog #324230, Corning) was tested following the protocol as recommended by the manufacturer, diluted 1:30 in cell media at 4 °C and 100 μL of diluted solution were added per well in the 96 well plate and incubated 1.5 h at room temperature or at 4 °C overnight before cell seeding onto the gel.

All cell lines and PBMC were incubated for 24 h from seeding to fixation. The following cell numbers per well was used throughout the study: 75,000 Jurkat cells, 25,000 THP1 cells, 12,000 HEL cells, 75,000 K-562 cells, 150,000 REH cells and 8,000 U-2 OS cells. For PBMC 75,000 cells per well was seeded directly after isolation.

### PBMC and platelet isolation and seeding

PBMCs were isolated from freshly drawn blood using the BD Vacutainer® CPT™ Mononuclear Cell Preparation Tube (CPT) (BD Biosciences, Franklin Lakes, USA, 362761) with Ficoll-Paque as a density medium and sodium citrate as anticoagulant according to product manual guidelines.

Briefly, the CPT was filled with 7-8 ml of fresh blood and immediately centrifuged for 20 minutes at 1500 RCF. Afterwards approximately half of the plasma layer was aspirated and the cell layer with the mononuclear cells and platelets was collected in a new 15 ml centrifugation tube. The tube was then filled up to 15 ml with 1x phosphate buffered saline (1x PBS) (137 mM NaCl, 2.7 mM KCl, 10 mM Na_2_HPO_4_, 1.8 mM KH_2_PO_4_, pH 7.2) and the cells were mixed by inverting the tube 5 times followed by a centrifugation step for 15 minutes at 300 RCF. The supernatant was aspirated and the cells were resuspended in 1 ml 1x PBS by gently vortexing. The tube was then filled up to 10 ml with 1x PBS, the cells were mixed cells and centrifuged for 10 minutes at 300 RCF with brake on. Afterwards the cells were resuspended in 1 ml 1x PBS by gently vortexing. To remove remaining red blood cells the eBioscience 10 X RBC Lysis Buffer (Thermo Fisher Scientific, Waltham, USA, 00-4300-54) was diluted one to ten (1:10) in MilliQ water and 10 ml were added to the resuspended cells. The cells were incubated for 15 minutes at room temperature (RT) followed by a centrifugation step for 5 minutes at 300 RCF. The cells were then resuspended in PBMCs media (RPMI 1640 supplemented with 10 % fetal bovine serum (FBS), 1 % sodium-pyruvate, 1 % penicillin-streptomycin and 1 % L-glutamine) and a cell count with a Neubauer counting chamber was performed prior to seeding into the 96 well plates for immunofluorescence.

Platelets were isolated and prepared as previously described (13). Briefly, human venous blood was drawn into sodium citrate from healthy adult volunteers in accordance with an Oregon Health & Science University IRB-approved protocol. Platelet rich plasma (PRP) was prepared by centrifugation of anticoagulated blood at 200 RCF for 10 minutes. Platelets were further purified from PRP by centrifugation at 1000 RCF in the presence of prostacyclin (PGI_2_). Following wash with HEPES/Tyrode buffer (129 mM NaCl, 0.34 mM Na_2_HPO_4_, 2.9 mM KCl, 12 mM NaHCO_3_, 20 mM HEPES, 5 mM glucose, 1 mM MgCl_2_; pH 7.3), platelets were resuspended in HEPES/Tyrode buffer for sample preparation.

### Cell fixation and immunostaining

The IF sample preparation protocol used was based on the protocol established for adherent cells by Stadler et al (14). For the coating experiments, all steps were performed manually to Shortly, cells were fixed with ice cold 4 % paraformaldehyde (PFA)(43368.9M, VWR Stockholm Sweden) and for 15 min and permeabilized with 0.1 % (w/v) Triton X-100 solution diluted in PBS for 3 times 5 minutes. Primary antibodies were diluted in blocking buffer (PBS with 10 % FBS (F7524, Sigma-Aldrich) overnight at 4 °C. Following four washing steps with 1x PBS, secondary antibodies were incubated at RT for 90 minutes, followed by staining with the nuclear stain 4’,6-diamidino-2- phenylindole (DAPI) at 2.28 μM solution for 10 minutes at RT. Eventually cells were washed four times with 1x PBS and then the entire well was filled with glycerol (85 %) as a mounting media before plates were sealed with aluminium foil covers.

For imaging of platelets, glass coverslips were coated with 50 μg/ml human fibrinogen in PBS for 1 hour prior to washing in PBS. Platelets (2×10^7^/ml) were incubated on fibrinogen-coated cover glass (45 min, 37 °C) prior to fixation and staining as previously described (15).

### Antibodies

Primary rabbit polyclonal antibodies generated within the Human Protein Atlas project and distributed by Atlas Antibodies (Stockholm, Sweden) were diluted to 2-4 μg/ml, a mouse monoclonal anti-alpha tubulin antibody (Ab7291, Abcam Sweden) diluted to 0,5 μg/ml) and a rat monoclonal anti-KDEL antibody (ab50601, Abcam Sweden) diluted to 2.5 μg/ml were used as common markers for the cytoskeleton and endoplasmic reticulum (ER) respectively. Secondary antibodies goat anti-rabbit IgG Alexa-Fluor 488 (**A11034)**, goat anti-mouse IgG Alexa-Fluor 555 (**A21424)** and goat anti-rat IgG Alexa-Fluor 647 (**A21247**) all from Thermo Fischer Scientific Sweden, were diluted to 2.5 μg/ml in blocking buffer and used to detect rabbit HPA antibodies, the anti-alpha tubulin antibody the anti-KDEL antibody respectively.

Adherent platelets were stained with antisera against vinculin (Sigma, V9131), filamin A (Santa Cruz Biotechnology, sc-17749), non-muscle myosin IIa heavy chain (MYH9, Abcam, ab89837), P-selectin (sc-8419), reticulon-4 (RTN4, sc-271878). Following staining with primary antibodies, platelets were stained with secondary antibodies and TRITC-phalloidin (Sigma, P1951) (15).

### Automated sample preparation protocol

The liquid handling robot Tecan Evo Freedom (Tecan Nordic Ab, Stockholm Sweden) was used for automated plate preparations of cells and three new scripts were created and tested on the panel of different suspension cell lines. Each of the new scripts were optimized to change one off the following parameters during the sample preparation: 1) the aspiration volume in all steps was reduced to always leave liquid at the bottom of the plates, 2) the number of washing steps was reduced or 3) the pipetting speed was reduced to lowest possible. All scripts used by the Tecan Evo Freedom can be found in the repository at Github, DOI 10.5281/zenodo.3463285(16).

### Image acquisition

For evaluation of cell distribution whole wells were acquired automatically with the Leica inverted DMi8 widefield microscope (equipped with a HC PL FLUOTAR 4x/0.13 or 20x/0.4 N/A air objective and Hamamatsu Flash 4.0 V3 camera package) using the Leica Application Suite X (LAS X) software with the Leica LASX Navigator function.

The Leica SP5 or SP8 confocal laser scanning microscope (CLSM), equipped with a 63x/1.4 N/A oil immersion objective, was used to acquire high resolution images to allow for quality evaluation of fixation and organelle staining patterns. The images were acquired at RT in sequential steps using the following settings: 16-bit acquisition, pixel size 0.08 μm x 0.08 μm, line averaging of 2 and a pinhole of 1 airy unit (AU), scan speed 600.

### Image analysis and cell counting

Images acquired from the DMI8 microscope for quantification and cell distribution purposes were processed using Fiji Is Just ImageJ (Fiji) (17) to enhance contrast of the cell nuclei for illustrative purposes. For high resolution images brightness and contrast was automatically adjusted where needed for illustration purposes using Fiji. Cell Profiler (18) was used to quantify the number of cells in each well from the different coatings and automated sample preparation experiments. For each cell line 16 replicate samples (wells) were prepared (7 wells for REH due to insufficient number of cells for 16 wells). For every sample, image analysis was used to count cell nuclei and calculate the fraction of remaining cells compared to seeded cells.

## Results

### Evaluation of suspension cell adherence to a variety of coatings

As a first step in developing a high throughput protocol for immunofluorescence of suspension cells, different coating materials were used to find the coating with the best prerequisites for retaining cells at a good density and even distribution across the sample well. To ensure as good image quality as possible, only glass bottom plates were used. These initial experiments focused on the human T-cell derived Jurkat cell line which represents a commonly used suspension cell line model in immunology and hematology studies. Replicate wells were coated with the different ECM proteins fibronectin (FN), laminin (LN), or Matrigel, containing a mixture of LN, collagen and entactin (19). The synthetic polymers Poly-L-ornitin and poly-L-lysine were also evaluated as they are popular coatings for plastic culture vessels and considerably cheaper than the biological coatings.

The highest number of remaining cells after the complete IF protocol was observed with fibronectin. These samples show an even distribution of cells in the well in a reproducible way (**Fig 1a**). For all other coatings, the overall cell density was lower even with the highest concentration of coating, and varied between different regions of the same well. In many cases only small islands of cells or cells at the immediate edges of the well was observed, as represented in (**Fig 1b.)** For Matrigel the results between the wells varied significantly, both between the two coating protocols used (incubation for a short time at RT vs long time at 4 C°) and for the replicate wells. The cells also clumped together on this surface resulting in a very uneven distribution of cells. The Matrigel protocol was also the most challenging coating protocol to use, requiring fast handling of the material with chilled pipette tips. **Supplementary Figure 1** show a subset of all wells acquired for each of the five different coatings and concentrations.

**Figure 1.**
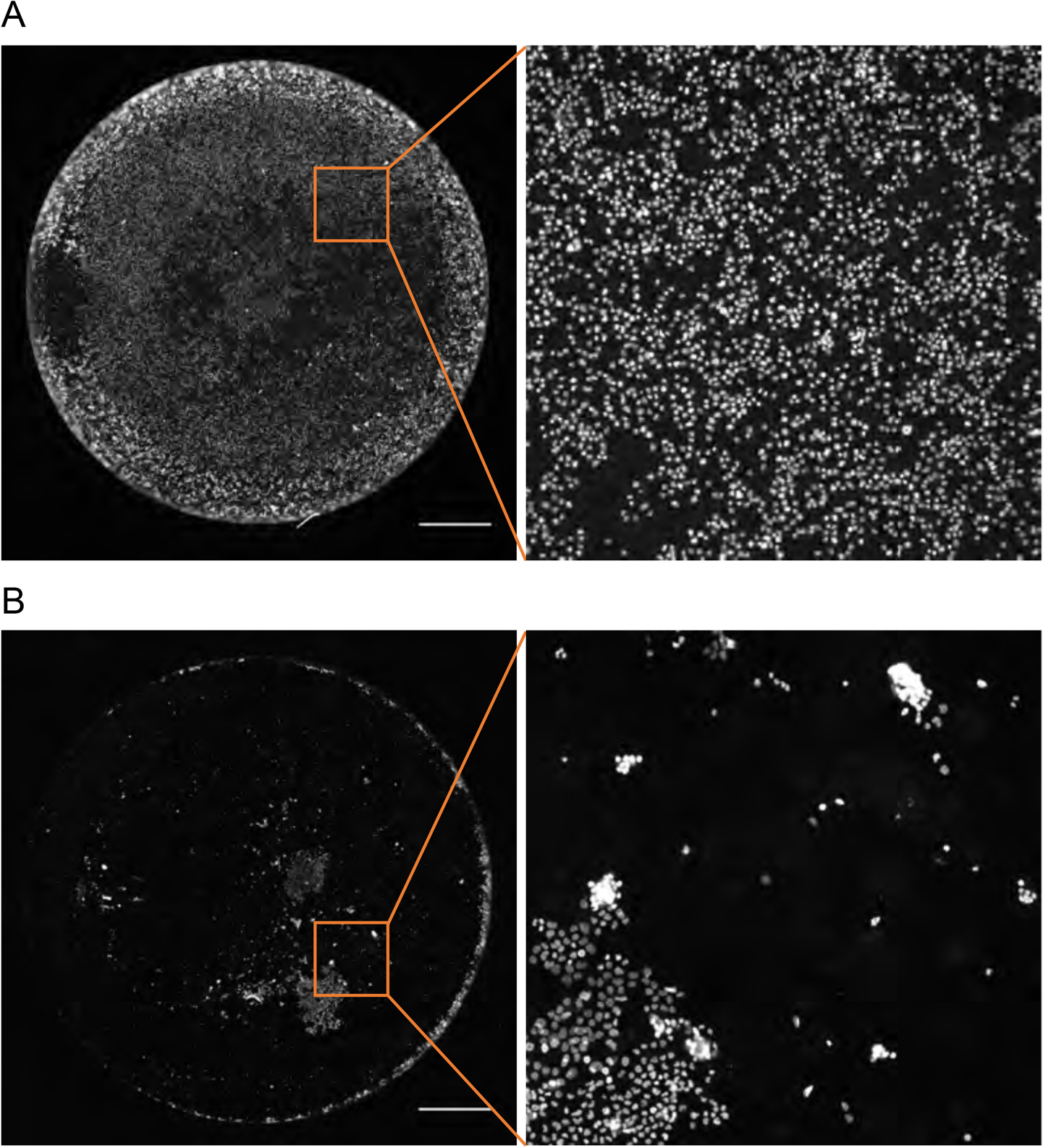
Representation of differences in cell density and distribution in two wells coated with fibronectin (**A)** and poly-L-lysin (**B**) from a 96 well plate. Cell nuclei are stained with Dapi and wells acquired with a 20x air objective. Scale bar is 1000 μm and cropped regions equals 1000×1000 μm.

As we aimed to obtain a reproducible protocol suitable for high throughput sample preparation and imaging, we selected FN for coating throughout the rest of the study.

### Automated sample preparation of suspension cells using liquid handling robotics

Next, to standardize and ensure the reproducibility of high throughput sample preparation of suspension cells, we tested a previously developed automated liquid handling protocol with a Tecan Evo Freedom “robot”, on five different suspension cell lines: HEL, THP1, REH, K-562 and Jurkat. To compare the fraction of remaining cells to a “best case scenario” the adherent cell line U-2 OS was used as a reference.

**Fig 2a** shows the distribution of cells in a representative well for each cell line. The images show that HEL, THP1 and U-2 OS to a large extent remain in the wells after completed sample preparation while most of K-562, Jurkat and REH cells were detached using this protocol. Quantitative data from cell nuclei counting reveal that for both HEL and THP1 the number of detected cells exceeds that of the seeded cell number, showing that cells are not only remaining but also divide from time of seeding until fixation 24 h after (**Supplementary Figure 2**). Further, the high number of cells and even distribution of HEL, THP1 and U-2 OS cells was observed for all wells analyzed, with a coefficient of variation (CV) of 7 % for HEL, 19 % for THP1, and 14 % for U-2 OS. For K-562, Jurkat and REH many cells were lost and only a small fraction of cells (19, 17 and 13 % respectively) remained, mostly at the edges of the wells. The remaining cell number for these cell lines also varied more, with a CV of 46 % for K-562, 23 % for Jurkat and 16 % for REH. The CV for REH was better than for many cell lines, however as a consequence of having almost no cells left in any of the wells.

**Figure 2.**
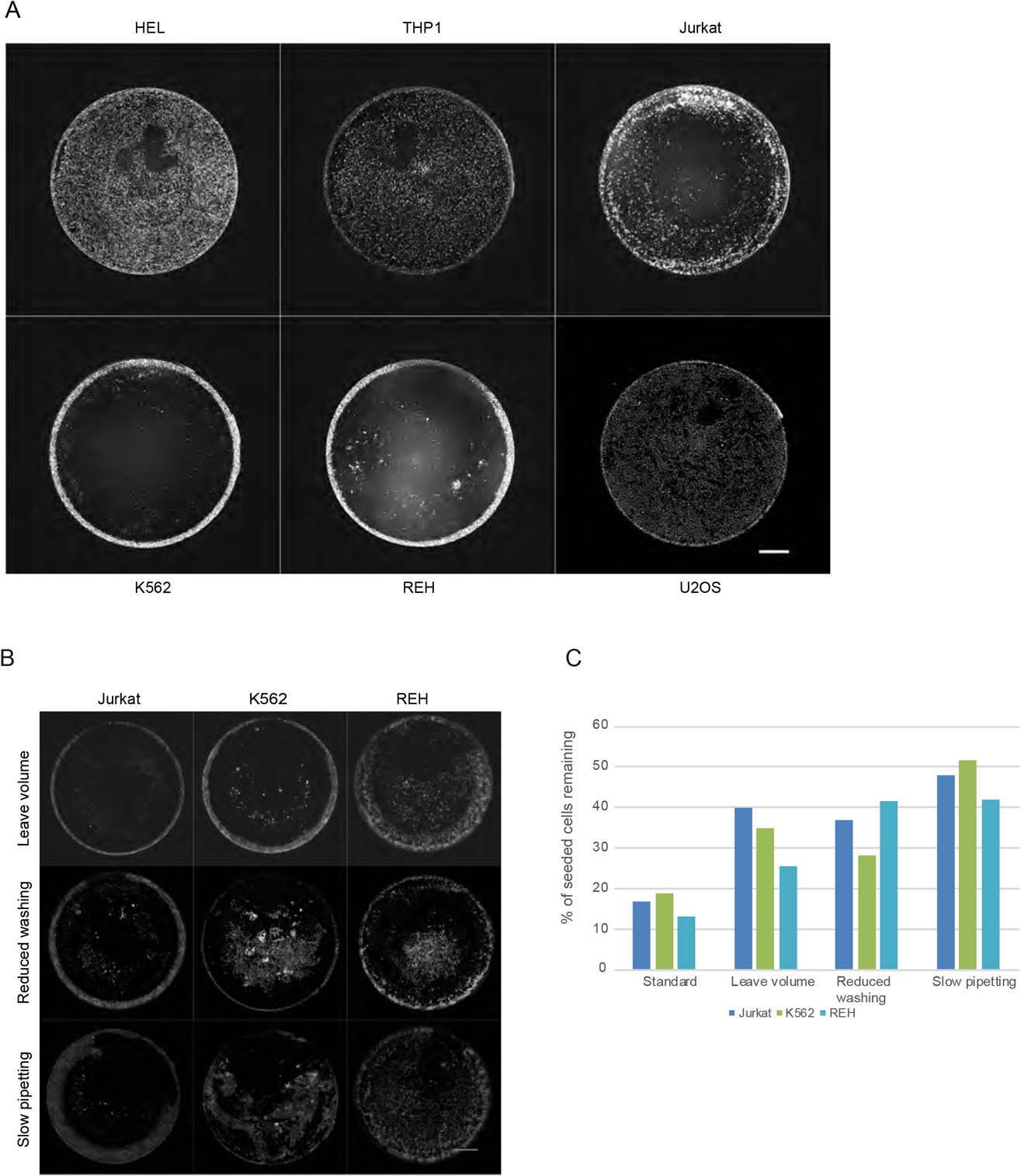
**A.** Representative images showing cell distribution for each cell line using the automated protocol normally used for adherent cells. Images were acquired with a 20x air objective. *Upper row*: HEL, THP1 and Jurkat. *Lower row:* K-562, REH and U-2 OS. Scale bar: 1000 μm. **B.** Representative images showing cell distribution for K-562(left), REH (middle) and Jurkat (right) cell line using the three optimized protocols: Leave volume (upper row), reduced washing steps (middle row) **C:** Bar chart showing the average fraction of remaining cells from replicate wells for K-562, REH and Jurkat cells, for each of the three optimized automated protocols.

To improve the fraction of remaining K-562, Jurkat and REH cells the pipetting protocol for the liquid handling robot was optimized. While the conditions for fixation, permeabilization and immunostainings was identical to the original protocol, the number of washing steps or other instrument parameters were optimized. In total three different protocols were tested differing from the standard protocol as follows: 1) The number of washing steps after primary and secondary antibody staining were reduced from four to three 2) The aspirated volume was decreased to always leave some liquid in the wells and 3) The pipetting speed was reduced from 4 to 0.5 μL/s. 16 samples of each cell line (8 for REH) were prepared, imaged and evaluated by comparing remaining cells after completed sample preparation with the standard protocol and U-2 OS reference cell line**. Figure 2b** shows representative wells from the different pipetting protocols for Jurkat, K-562 and REH and the average number of cells remaining from the different protocols (**Fig 2c**). Based on this, the slow pipetting protocol resulted in the largest fraction of remaining cells for all cell lines. Compared to the standard protocol the average fraction of remaining cells increased from 17 to 48 % for Jurkat, 19 to 52 % for K-562, from and 13 to 42 % for REH. The coefficient of variation (CV) was also calculated for the respective cell line and compared to the original standard protocol. For Jurkat the CV was the same, whereas it was reduced from 46 to 32 % for K-562. While the overall number of remaining REH cells was greatly improved, the CV increased from 16 to 33 %. These results suggest that lower pipetting speed gives better prerequisites for retaining a larger fraction of cells during the sample preparation, although other parameters such as cell seeding can influence the result of individual wells. The quantitative data for each cell line and protocol is shown in **Supplementary Figure 3.**

Compared to HEL and THP1, the Jurkat, K-562 and REH cells detached to a much higher extent also with the optimized protocol. To investigate this further, we looked at RNA expression data (normalized transcripts per million reads – tpm) generated within the Human protein Atlas of all major cell adhesion proteins in the integrin and cadherin families. When comparing the sum of RNA transcript levels for these genes, a lower total expression of cell adhesion proteins was seen for K-562 and REH compared to HEL and THP1. In fact, the HEL and THP1 has expression levels similar to the adherent cell line U-2 OS **(Table 2).** As fibronectin was used as coating material, the expression levels of the integrins ITGAV and ITGB1 should be the most important as these are the integrin subunits with specific affinity for fibronectin [15, 18). Although the expression of these integrins is low in all cell lines as compared to U-2 OS, we speculate that the higher abundance of cell adhesion proteins overall in HEL and THP1 is favorable for improved attachment to any surface.

### Manual and automated sample preparation of freshly isolated PBMC and platelets

To test the optimized slow pipetting protocol in primary cells, cells isolated from freshly withdrawn blood (PBMC) were seeded and prepared either manually or using the optimized automated protocol with slow pipetting. **Figure 3** show a representative whole well image and cropped region of cell distribution from the respective preparations (Fig 3a) and the number of remaining cells in individual wells from manual and automatically prepared samples (Fig 3b). The fraction of remaining cells is comparable for the manual and automated sample preparation, with 14 % for both manually and automatically prepared wells and a CV of 36 and 28 % respectively. Even though this fraction is low, it shows that the automated protocol can be used as successfully as the manual one. Further, the true fraction of remaining cells is likely higher than 14 %, as platelets that are part of the PBMC sample when counting cells prior to seeding, cannot be detected as they lack a cell nucleus and thereby contains no Dapi stain.

**Figure 3.**
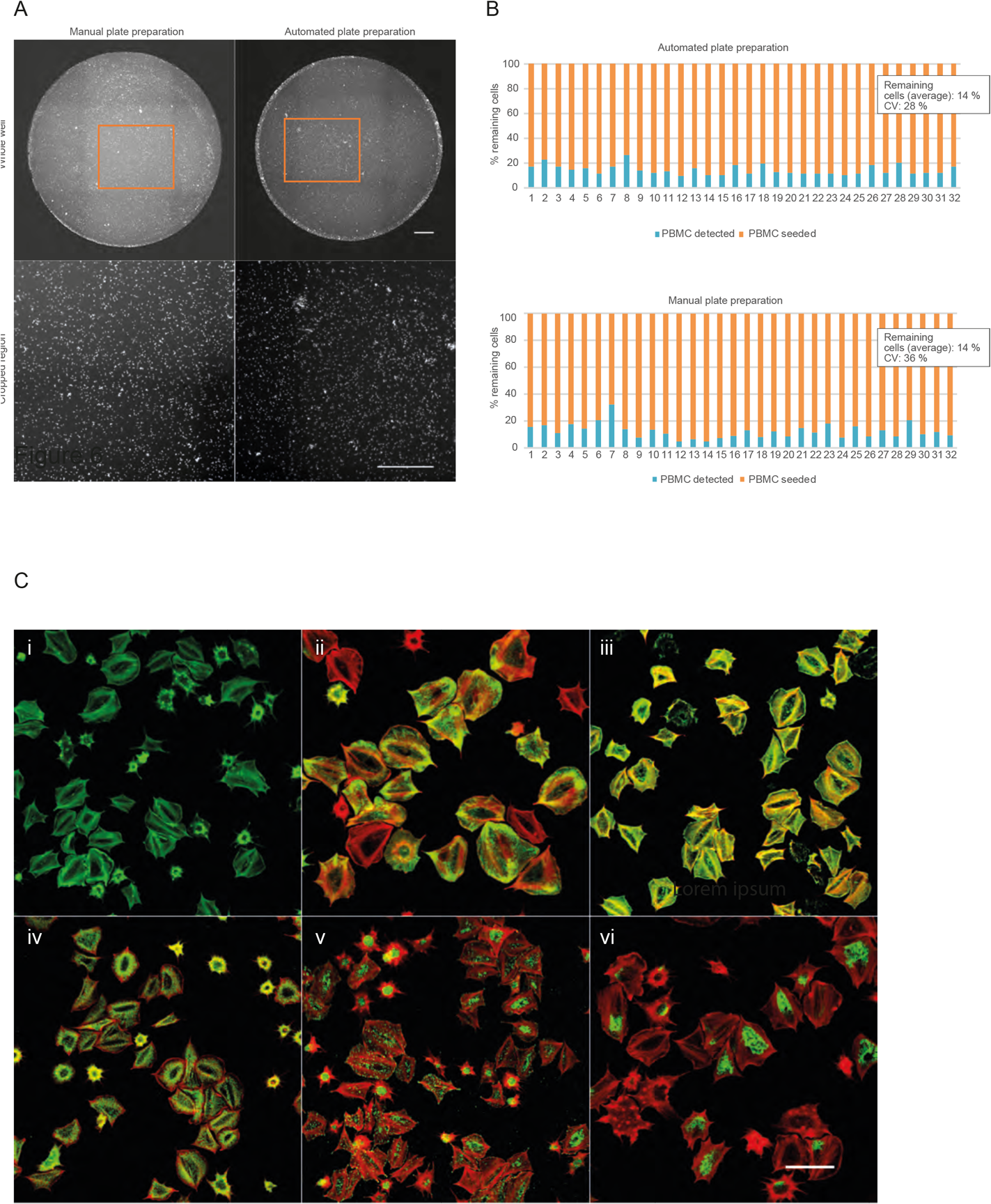
**A:** Representative images of a whole well and corresponding cropped region from a manual versus automatic plate preparation. Scale bar is 1000 μm and cropped regions equals 1000×1000 μm. **B:** Bar chart showing fraction of remaining PBMC in 32 individual wells from a manual versus automatic plate preparation. **C:** Confocal images of selected structures in platelets: F-actin stained with phalloidin (A), actin filaments/focal adhesions stained for vinculin (B), filamin A (C) and myosin heavy chain 9 (D), alpha granules stained for P selectin (E) and endoplasmic reticulum visualized by staining of RTN4. Red stain (were present) represents microtubules. Images are acquired with a 63x oil immersion objective Scale bar: 10 μm.

To evaluate the quality and overall condition of blood cell preparations after fixation and sample processing, platelets were used as a representative peripheral blood cell for additional imaging analyses. Following fixation and permeabilization, washed human platelets adherent to fibrinogen were stained for cytoskeletal structures and other marker proteins to visualize and evaluate their morphology (Fig 3c). In addition to staining actin filaments with phalloidin as a marker of F-actin, the cytoskeletal protein vinculin (VCL), the focal adhesion protein filamin A (FLNA), the non-muscle myosin IIa heavy chain (MYH9), the platelet alpha granule marker P-selectin (SELP, CD62P) and the ER marker RTN4 were stained (20). Immunofluorescence microscopy analysis confirmed the presence of intact cytoskeletal and other intracellular structures representative of adherent platelets.

### High resolution imaging of organelle markers in suspension cells

Finally, Jurkat cells were stained and imaged at high resolution to evaluate fixation quality and whether different subcellular structures could be distinguished within these suspension cells. **Figure 4** shows representative cells stained for a few different organelle marker proteins. For nuclear targets SRRM2 (nuclear speckles) and DNAJB2 (nuclear membrane) the staining patterns are very similar to those of adherent cells, and thus easily annotated. The same applies for staining of the plasma membrane, endoplasmic reticulum and cytoskeletal structures like intermediate filaments, here represented by staining of PEBP1, CALR and PGM2 respectively. Also the Golgi apparatus here stained for GOLGA2, is easy to identify given it’s dense structure and location in close proximity to the nucleus. Other cytosolic organelles such as mitochondria (here represented by staining of ETFA) and vesicles (here represented by peroxisomes stained for AGP5) can also be reliably identified. These results suggest that the optimized protocol in combination with confocal microscopy is suitable for automated sample preparation for subcellular profiling in suspension cells, even though these cells display a small cytosol compared to most adherent cells types.

**Figure 4.**
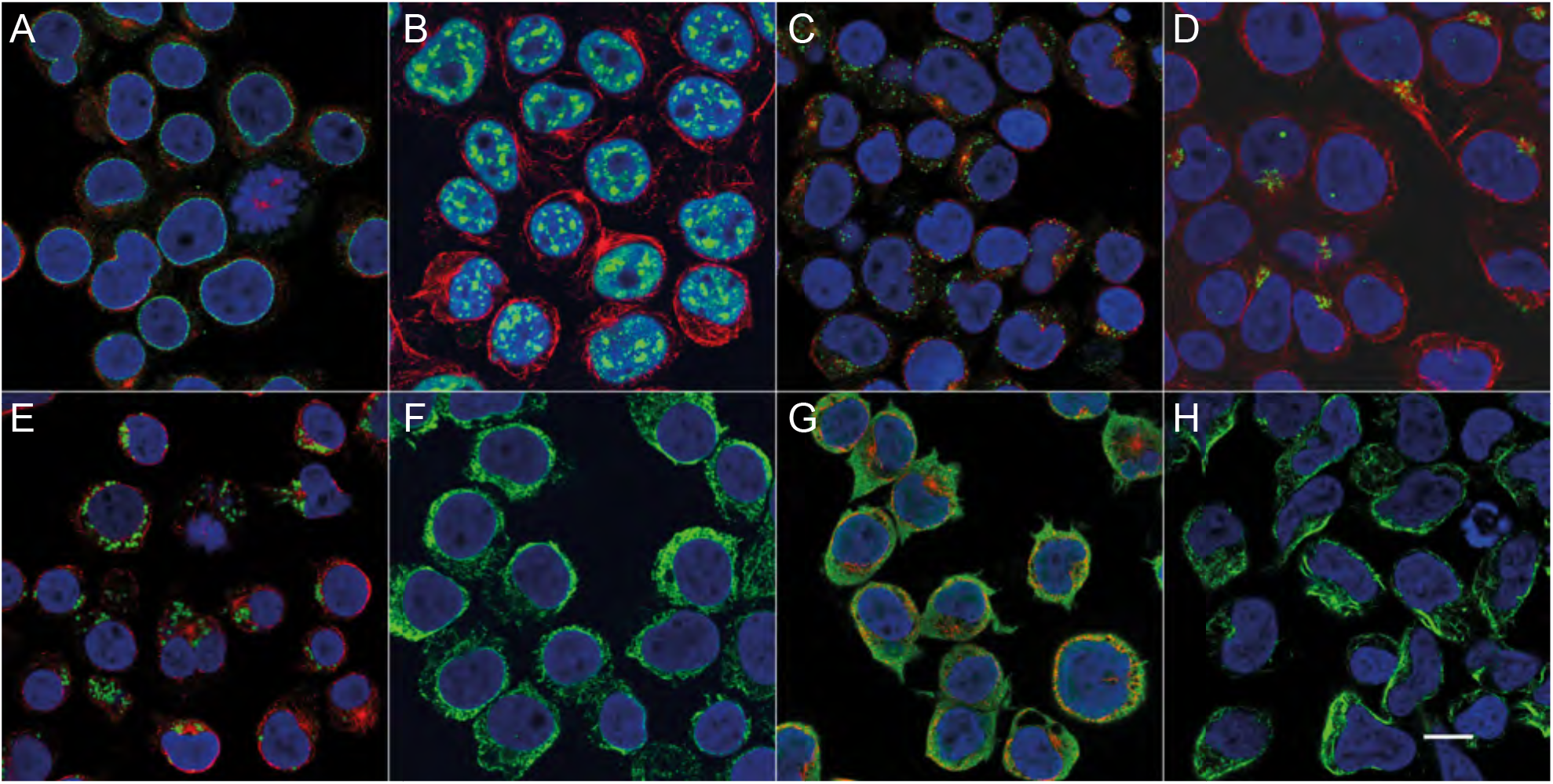
Confocal images of selected organelles in Jurkat cells: nuclear membrane (A), nuclear speckles (B), peroxisomes (C), Golgi apparatus (D), mitochondria (E), endoplasmic reticulum (F), plasma membrane (G) and intermediate filaments (H). All cells are counterstained with Dapi (blue). Red (were present) represents microtubules. Images are acquired with a 63x oil immersion objective Scale bar: 10 μm.

## Discussion

In this work, we optimized a protocol for sample preparation for immunofluorescence of suspension cells, compatible with cell lines as well as primary PBMC and platelet cells. In addition, this protocol was automated using liquid handling robotics to allow for high-throughput applications.

By optimizing different parameters, the fraction of remaining cells could be increased for all tested cell lines, with the best results obtained by reducing the pipetting speed to minimal. This parameter is in our experience the most critical for a successful manual sample preparation of any cell line, as it reduces the shear forces on the cells. Given that all individual parameters (reduced aspiration volume, reduced number of washing steps and reduced pipetting speed) resulted in a larger fraction of remaining cells, it is likely that a protocol combing all of these parameters would generate even better results in terms of remaining cells. However, the reduced number of washing steps is not ideal since it might affect background and thus overall quality of the images. In case of the parameter of a lower volume being aspirated throughout the protocol, this slightly changes the concentration and exposure time of reagents and should therefore be evaluated more carefully before being implemented. Still, this is something to consider for further improvements.

We observed different overall success in remaining cell number and CV for the different cell lines used in this study. Our evaluation of cell adhesion protein expression by RNA sequencing data, showed higher expression of adhesion proteins in HEL and THP1, lower for Jurkat and lowest for K-562 and REH. This correlates well with the number of remaining cells and also with the robustness, as HEL and THP1 where the cell lines with highest fraction of reaming cells and smallest CV. Based on this, the RNA expression data available in the HPA database for a broad panel of cell lines, could be a useful tool when selecting cell lines for imaging, or in selecting the most suitable coating for a particular cell line, depending on the expression of specific integrins. Besides expression of cell adhesion proteins, the size of cells could be relevant for how well they attach. For example, the HEL and THP1 cell lines are considerably larger than K-562 and REH, suggesting more binding events to the surface and thus a stronger attachment. In our experience, smaller cells like K-562 and REH more easily aggregates and are harder to homogenize in the solution prior to seeding. We believe this can have an impact of the overall CV of remaining cells as cell number within wells are more likely to differ than those of larger cells. To get a more precise fraction of remaining cells in each well, cells would have to be counted before starting the sample preparation. In this work however, the primary aim was to increase the number of remaining cells by optimizing sample preparation, excluding cell seeding conditions.

In addition to establish an automated protocol for sample preparation allowing for high throughput, we also demonstrate how immunofluorescence can successfully be used for protein localization at subcellular level in suspension cell lines as well as in platelets. This enables systematic exploration of the intracellular landscape of different cell types derived from blood, and provide an important complement to flow-based methods lacking the spatial information obtained from imaging. Recently generated data within the framework of the HPA using Mass Cytometry (CyTOF) and RNA sequencing of 18 specific cell types from PBMC, suggest almost 1500 protein coding genes to have an elevated expression in any of the six hematopoietic cell lineages (T-cells, B-cells, NK cells, monocytes, granulocytes and dendritic cells). This makes it highly relevant to better explore their intracellular proteomes. The data above is part of the HPA effort in creating a Blood Atlas, which was recently released as a complement to the three existing sub atlases; Tissue Atlas, Cell Atlas and the Pathology Atlas.

The optimized protocol from this work, allows for high throughput sample preparation and imaging of suspension cells, that in combination with the proteome wide collection of antibodies generated within the Human Protein Atlas project presents opportunities to create a subcellular map of immune cells as a way to better understand the immune system in health and disease.

## Acknowledgements

We acknowledge Knut and Alice Wallenberg foundation (KAW) for their generous support to the Human Protein Atlas project, which instrumentation has been used to conduct this work and Science for life laboratories for funding the Cell Profiling Facility at Royal Institute of Technology that has been instrumental for the entire work from cell cultivation to sample preparation and microscopy.

---

**Supplementary Figure 1.**
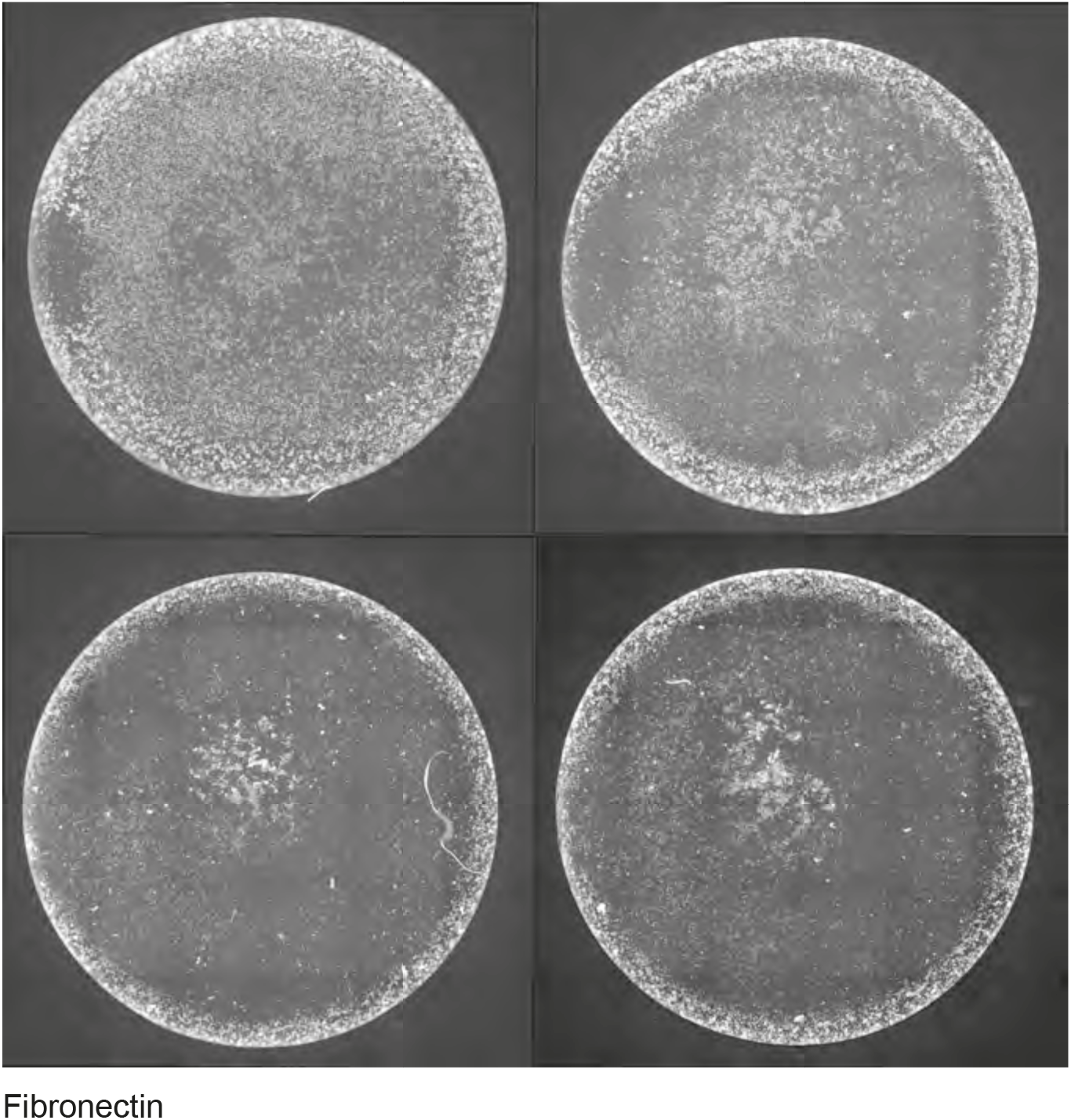
Images showing distribution of Jurkat cells in wells coated with five different coatings at various concentrations. A. Four replicate wells coated with 12.5 μg/ml fibronectin. B. Representative wells coated with Poly-l-ornithin, poly-l-lysine and laminin with a concentration of 12.5 25, 50 or 100 μg/ml. C. Representative wells coated with Matrigel diluted 1:30 and incubated 1.5 h at RT or overnight at 4 °C.

**Supplementary Figure 2.**
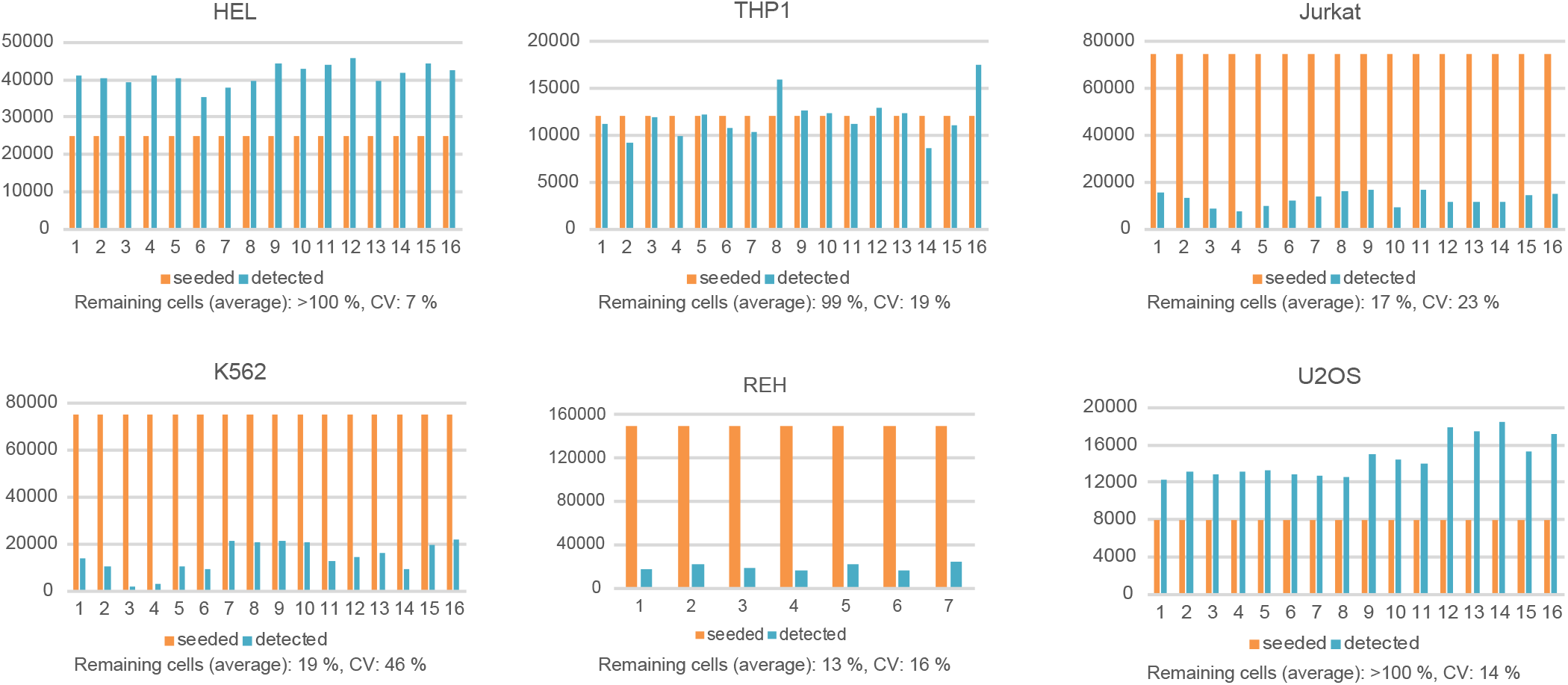
Bar charts for each of the six cell lines showing the number of seeded (orange) and remaining (blue) cells (vertical axis) using the automated standard protocol. Each chart display values for 16 or 8 replicate wells (horizontal axis) along with a calculated CV and average number of remaining cells.

**Supplementary Figure 3.**
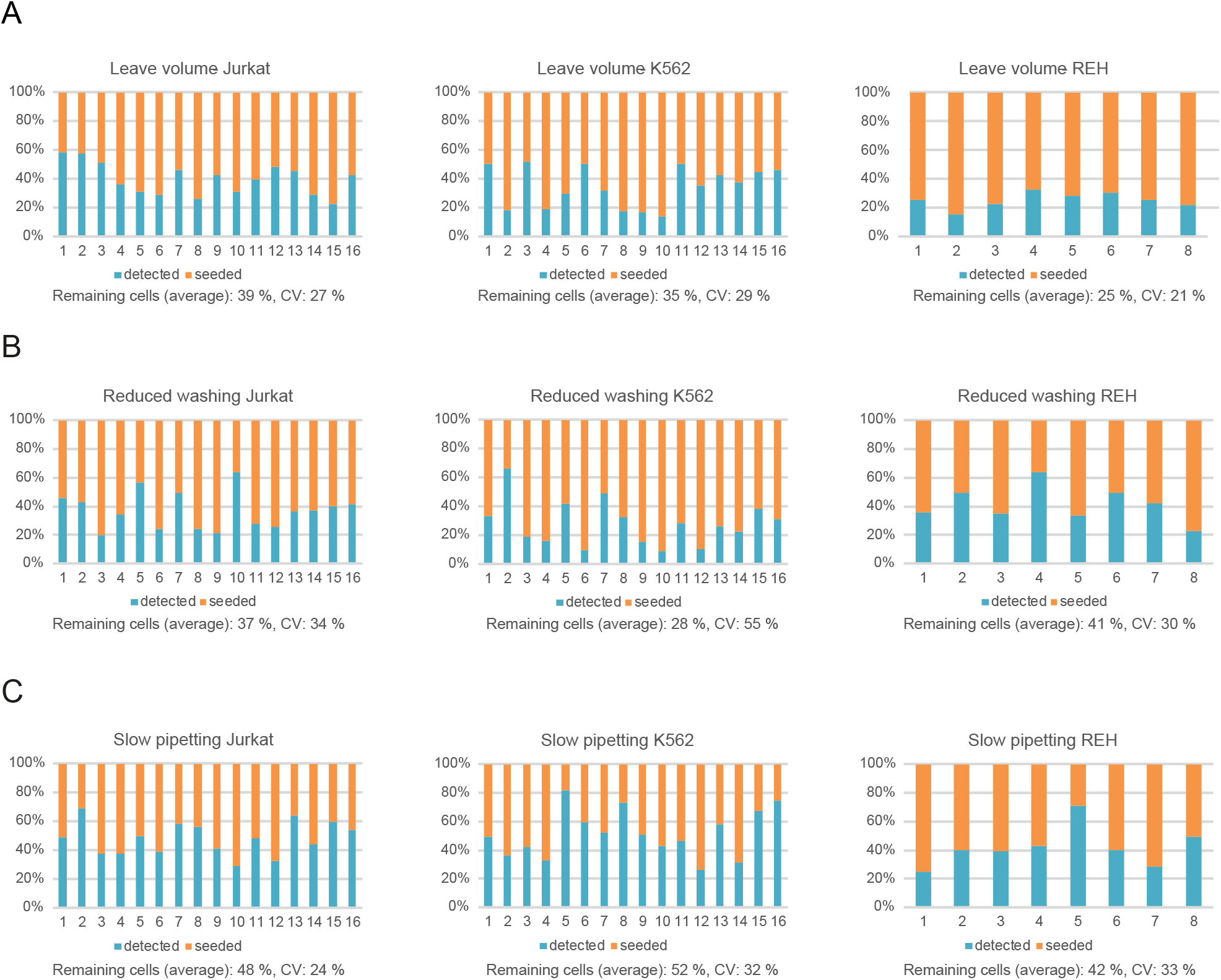
Bar charts for Jurkat, K-562 and REH cell lines showing the number of seeded (orange) and remaining (blue) cells (vertical axis) using the **A.** leave volume protocol **B** Reduced washing protocol and **C** Slow pipetting protocol. Each chart display values for 16 or 8 replicate wells (horizontal axis) along with a calculated CV and average number of remaining cells.

